# An automated workflow for quantifying the formation of synuclein aggregates in human dopaminergic neurons

**DOI:** 10.64898/2025.12.19.695578

**Authors:** Chanshuai Han, Emmanuelle Nguyen-Renou, Faiza Benaliouad, Wen Luo, Carol X-Q Chen, Aeshah Alluli, Lorenza Villegas, Lenore K Beitel, Irina Shlaifer, Wolfgang E Reintsch, Andrea I Krahn, Esther Del Cid Pellitero, Edward A Fon, Thomas M Durcan

**Affiliations:** The Neuro’s Early Drug Discovery Unit (EDDU), McGill University, 3801 University Street, Montreal, Quebec H3A 2B4, Canada; Neurodegenerative Disorders Research Group, Department of Neurology and Neurosurgery, Montreal Neurological Institute-Hospital (The Neuro), McGill University, 3801 University Street, Montreal, QC H3A 2B4, Canada

**Keywords:** α-syn: alpha-synuclein, DA NPCs : dopaminergic neural progenitor cells, DNs: dopaminergic neurons, iPSCs: induced pluripotent stem cells, PD: Parkinson’s disease, PFFs: preformed fibrils, pS129-syn: phosphorylated alpha-synuclein at serine 129

## Abstract

Parkinson’s disease (PD) is a neurodegenerative disorder characterized by alpha-synuclein (α-syn) aggregates termed Lewy bodies. To model PD pathology *in vitro*, preformed fibrils of α-syn (PFFs), which can be taken up by cells, provide a seed that drives misfolding and aggregation of endogenous α-syn, with new aggregates amplifying this process. External application of PFFs to dopaminergic neurons (DNs) increases aggregate formation, marked by α-syn phosphorylation at serine 129 (pS129-syn), a pathological PD marker. Building on this, we developed an automated synuclein seeding assay to quantify new α-syn aggregates in iPSC-derived DNs. Using pS129-syn as a readout, we show that PFFs elicit a time- and dose-dependent increase in pS129-syn aggregates. Our high-throughput assay further revealed that aggregate formation depends on endogenous α-syn levels. Treatment with PFFs produced a greater increase in pS129-syn aggregates in iPSC DNs derived from a PD patient with a triplication in the *SNCA* gene, which encodes the α-syn protein and which elevates total α-syn levels, relative to DNs from an isogenic iPSC line from the same individual, in which the *SNCA* gene mutation had been corrected by CRISPR/Cas9. In contrast, no pS129-syn signal was detected in neurons in which all copies of the *SNCA* gene had been knocked out (KO). This high-content imaging assay for synuclein seeding offers a platform for assessing compounds and therapeutics that may impede α-syn aggregate formation.

## 1. INTRODUCTION

Parkinson’s disease (PD) is a neurodegenerative disorder characterized by a progressive loss of dopaminergic neurons (DNs) and an accumulation of proteinaceous inclusions, termed Lewy bodies (LBs), in which alpha-synuclein (α-syn) is the protein present in highest abundance [1, 2]. Under physiological conditions, α-syn is a neuronal protein enriched in presynaptic terminals [1], although its physiological function details remain unclear. However, under pathological conditions, α-syn can misfold, forming oligomers and small fibrillar structures. These structures can subsequently recruit endogenous α-syn to form intracellular aggregates or LBs in neural soma and Lewy neurites (LNs) in axons and dendrites, coupled with the recruitment of membranous organelles, multivesicular bodies and lysosomes into these inclusions [2–4]. Posttranslational modifications (PTMs) of α-syn, particularly phosphorylation at serine 129 (pS129-syn) is identified as a key pathological hallmark of these LBs observed in patients with sporadic or genetic forms of PD [5]. More recently, PTMs on soluble α-syn have been shown to modulate the interaction of soluble α-syn with pathological α-syn to amplify the aggregation transmission process [6].

A number of cellular and animal models have been developed to study α-syn pathology in PD. Some of these models were based on the overexpression of wildtype or mutant α-syn alone [7], or coupled to treatment with other stress inducers [8]. However, the α-syn inclusions formed in these models were driven by overexpressing wild type α-syn or mutant α-syn (A53T and A30P), mutations commonly associated with the formation of α-syn aggregates. Recently, other types of models with α-syn preformed fibrils (PFFs) have started to attract attention. It has been shown in recent studies that the addition of exogenous α-syn PFFs can act as a seed, taken up into the cell by macropinocytosis [9], to initiate the misfolding and aggregation of endogenous α-syn in both cellular and animal models; ultimately forming inclusions that recapitulate the biochemical, morphological, and structural features of the LBs [4, 10–14]. Thus, the development of PFFs opened a number of avenues that could be explored for studying how synucleinopathies arise and for identifying therapies that can impede or reduce their spreading.

*SNCA*, the gene encoding the α-syn protein, was the first gene to be associated with familial Parkinson’s disease [15]. Missense mutations of *SNCA* cause an autosomal dominant form of familial Parkinson’s. Moreover, disease copy number variants of *SNCA* gene [16–20], that include duplication and triplication mutations, have shown a dose-dependent effect, where patients with a triplication of the gene display a more severe clinical phenotype, with an earlier onset than patients with the duplication mutation [21–24]. Using induced pluripotent stem cells (iPSCs) generated from a patient with a *SNCA* gene triplication (*SNCA* Tri), in a previous study [25] we showed an age-dependent increase in α-syn levels and aggregation in human midbrain organoids (hMOs) generated from the *SNCA* Tri line relative to control hMOs generated from an isogenic gene-corrected control iPSC line. Building on these earlier findings, we developed an α-syn seeding assay using the *SNCA* Tri iPSC from one PD patient, its isogenic control, where the SNCA mutant copies of the patient line have been deleted by CRISPR/Cas9 and its SNCA knock-out line, where all SNCA copies have been deleted by CRISPR/Cas9, to quantify the contribution of *SNCA* copy number and total α-syn levels on the capacity of α-syn PFFs to initiate α-syn aggregates. We observed that α-syn PFFs recruited endogenous α-syn to form LB-like inclusions in human DNs. We further adopted high-throughput imaging and analyzing workflows to develop a scalable screening assay monitoring the formation of α-syn aggregates in human DNs that could be used for identifying potential small molecules that could reduce the formation of α-syn aggregates.

## 2. MATERIALS AND METHODS

### 2.1. Cell-line information and ethical approval

The use of iPSCs and iPSCs derived cells in this research was approved by the McGill University Research Ethics Board (A03-M19-22A / eRAP 22-03-027). Human iPSC lines used in this manuscript are from a PD patient with a triplication of the *SNCA* gene locus (*SNCA* Tri), its isogenic control lines generated by CRISPR-mediated deletion of either two copies (Isog Ctrl) or all four 4 copies (*SNCA* KO) of the *SNCA* gene from the same parental *SNCA* trip line. Sanger sequencing of the *SNCA* region, *SNCA* copy number confirmation and iPSC quality control profiling was described previously [25].

### 2.2. Maintenance of iPSCs and dopaminergic neuron differentiation

*SNCA* Tri, Isog Ctrl and *SNCA* KO iPSC cultures were maintained as feeder-free cultures following the protocols described previously [25, 26]. iPSCs were plated onto Matrigel (Corning Millipore, Cat#354277)-coated plates containing mTeSR1 medium (Stemcell Technologies, Cat#85850). The culture medium was changed daily until cells reached ∼80% or required confluency (usually 5-7 days after plating). The cells were next either passaged, frozen, or differentiated into cells of interest.

Ventral midbrain dopaminergic neural progenitor cells (DA NPCs) from these three lines were generated following a previously described monolayer method [27]. iPSCs were plated into Matrigel-coated T25 flasks containing DA induction medium ((DMEM/F12 (Thermofisher scientific, Cat#10565042) supplemented with N2 (Thermofisher scientific, Cat#,17502048) B27 (Thermofisher scientific, Cat#17504044) supplements and MEM NEAA (wisent, Cat#321-011-EL), BSA (1 mg/mL; Invitrogen, Cat#15260-037), Noggin (Peprotech, Cat#120-10C 200 ng/mL), SHH (200 ng/mL; Genscript, Cat#Z03067 C24II), CHIR-99021 (3 μM; Selleckchem, Cat#S2924), SB431542 (10 μM; Selleckchem, Cat#S1067) and FGF-8 (100 ng/mL; Peprotech, Cat#100-25)). Media was changed daily, and cells were dissociated with Gentle Cell Dissociation Reagent (STEMCELL Technologies, Cat#07174) and split in a 1:2 ratio into Matrigel coated T25 flasks on day 7 and day 14. On day 21, cultures were cryopreserved as DA NPCs in FBS (Thermofisher, Cat#12484028) containing 10% DMSO (Fisher, Cat#BP231-1).

The final differentiation of the DA NPCs into DNs was achieved as previously described [28]. DA NPCs were thawed from preserved frozen stocks and cultured in PO ( 10 μg/mL; Sigma, Cat#P3655)/Laminin (5 μg/mL; Sigma, Cat#L2020) coated T25 flasks with STEMdiff Neural Progenitor Basal Medium (STEMCELL Technologies, Cat#05834) supplemented with STEMdiff Neural Progenitor Supplement A (STEMCELL Technologies, Cat#05836), STEMdiff Neural Progenitor Supplement B (STEMCELL Technologies, Cat#05837) and Purmorphamine (2 μM; Sigma, Cat#SML-0868) for one week. After one-week recovery, DA NPCs were dissociated with StemPro Accutase Cell Dissociation Reagent (ThermoFisher, Cat# A1110501) into single-cell suspensions. 50,000 cells were plated onto PO (2 μM; Sigma, Cat#P3655)/Laminin (5 μg/mL; Invitrogen, Cat#23017-015) coated coverslips in 24-well plates; or 15,000 cells per well were plated into 96-well plates with dopaminergic neural differentiation medium (Brainphys Neuronal medium, STEMCELL Technologies, Cat# 05790) supplemented with N2A Supplement A (STEMCELL Technologies; Cat# 07152), NeuroCult SM1 Neuronal Supplement (STEMCELL Technologies; Cat# 05711), BDNF (20 ng/mL; Peprotech, Cat# 450-02), GDNF (20 ng/mL; Peprotech, Cat# 450-10), Compound E (0.1 mM; STEMCELL Technologies, Cat# 73954), db-cAMP (0.5 mM; Carbosynth, Cat# ND07996), Ascorbic acid (200 mM; Sigma Aldrich, Cat# A5960) and laminin (1 mg/mL, Sigma Aldrich, Cat# L2020).

### 2.3. Preformed fibril production and QC

α-syn preformed fibrils (PFFs) were generated from recombinant monomeric α-syn (Fig. S1A). Recombinant human α-syn was purified from a plasmid containing glutathione S-transferase (GST)–tagged full-length recombinant human α-syn expressed in BL21 (DE3) Escherichia coli. In parallel, a plasmid containing GST-tagged recombinant 3C enzyme was also expressed for cleaving the GST tag. The GST-tagged proteins were purified from the bacterial cell lysates by affinity column chromatography (GE Healthcare, Cat#17-5282-01) and the purified GST-α-syn protein was treated with GST-3C enzyme to remove the GST tag. Untagged α-syn protein was purified from the reaction using affinity column chromatography and further purified using size exclusion chromatography (GE Healthcare, Cat#28-9893-35) (Fig. S1B). Recombinant monomeric α-syn protein was then ready to be used for experiments directly, or for generating PFFs by inducing its aggregation. Untagged α-syn protein (monomeric form) was tested for bacterial endotoxin (kit from ThermoFisher Scientific, Cat#A39552) to ensure that levels were below <0.05 units of endotoxin/ml. Generation of α-syn preformed fibrils (PFFs) from recombinant monomeric α-syn was obtained after five days of shaking in a 37°C digital heating shaker dry bath (ThermoFisher Scientific, Cat#88880027) or thermomixer (Eppendorf, Cat#2231000574) at 1000 rpm. After shaking, α-synuclein preformed fibrils (PFFs) were sonicated using Bioruptor Pico sonicator (with the following settings: 50 cycles, 30 seconds ON/30 seconds OFF, 6°C water circulation). Aliquots of PFFs in 1.5-mL tubes were made and stored at −80°C.The characteristics of α-syn PFFs were validated prior to experimental use by performing quality control analyses of fibril length by EM (electron microscopy; Spirit (TEM), #FEI Tecnai G2 spirit Biotwin 120KV; Fig. S1C) and DLS (dynamic light scattering; Malvern Panalytical, # Zetasizer nano S, software 7.13; Fig. S1D). Only samples with a single peak with average size smaller than 100 nm are qualified for downstream experiments.

### 2.4. High-content Immunofluorescence (IF) imaging

Cells were washed with phosphate-buffered saline (PBS, Wisent Inc, Cat#311-010-CL), then fixed with 4% paraformaldehyde (ThermoFisher Scientific, Cat#28908) on coverslips or 96-well plates for 15 minutes. Samples were permeabilized with 0.2% TX-100 (Sigma-Aldrich, Cat#X-100) in PBS for 5 minutes and then blocked in PBS containing 0.02% Triton X-100 and 2% Bovine serum albumin (Invitrogen, Cat#15260-037) for one hour. Primary antibodies (Table S1) were diluted with PBS containing 0.02% Triton X-100 and 2% BSA and added to samples and incubated overnight. Samples were washed in PBS containing 0.02% Triton X-100 four times. PBS with 0.02% Triton X-100 and 2% BSA containing an appropriate dilution of secondary antibody (Table S1) was added to the samples and incubated for 60 minutes in the dark at room temperature. Samples were washed with PBS containing 0.02% Triton X-100 and nuclei were stained with Hoechst 333342 (Thermo Fisher Scientific, Cat#H3570). Images were acquired with Leica TCS SP8 confocal microscope or on an automated microscope high content system (six wells/condition; 25 fields/well) ThermoFisher Scientific CellInsight CX7 and analyzed with the Studio software.

### 2.5. RNA isolation, cDNA synthesis and qPCR analysis

RNA from iPSCs, DA NPCs and DNs was purified using the NucleoSpin RNA kit (Takara, Shiga, Japan, 740955) according to the manufacturer’s instructions. cDNA was generated using the iScript Reverse Transcription Supermix (BioRad, Hercules, CA, USA, 1708840). Quantitative real-time PCR was performed on the QuantStudio 5 Real-Time PCR System (Applied Biosystems, Waltham, MA, USA) using the primers listed in Supplementary Table S2. Raw data were processed using a custom Python script, available at https://github.com/neuroeddu/Auto-qPCR (accessed on 29 July 2020)[29, 30]. The cycle threshold (CT) values for technical triplicates were tested for outliers. Relative gene expression was calculated by using the comparative CT method (ΔΔCT method), where the endogenous controls were GADPH or ACTB expression. Expression of the genes of interest were calculated relatively to the endogenous control genes for each cell line.

### 2.6. Preformed fibril treatment and data analysis

DA NPCs were seeded into 96-well plates and maintained in DN maturation media for 2 weeks. After 2-weeks of neuronal maturation, DNs were treated with PFFs (at a final concentration of 500nM or at concentrations indicated for different experiments) or with the α-syn monomers (at the same concentrations as PFFs) and left 14 days in the presence of PFFs or the α-syn monomers. For the small-molecule compound tests, different compounds (Table S3) were added to DNs in addition to PFFs. After the PFF treatment, the DNs were fixed with PFA 4% and newly formed α-syn aggregates were visualized by IF staining with the phosphorylated α-syn antibody (Table S1). For samples subjected to phosphatase treatment, cells were fixed and permeabilized and then incubated with 50 units/μL Lambda protein phosphatase (New England Biolabs, cat P0753L) at 30°C for 8 hours before IF staining. Images were acquired in triplication (81 fields/well), except for the PFF conditions where 12 wells were acquired, on an automated microscope high content system (ThermoFisher Scientific CellInsight CX7 Pro High-Content Screening Platform) using a 20X objective and analyzed with Neuronal profiling BioApplication (HCS Studio 5.0 Cell Analysis Software, ThermoFisher Scientific). Nuclei were identified with Hoechst 33342 staining and were used as autofocus reference. Neurites were identified with TUBB3 staining through the neurite identification module and total neurite areas were set as Region of Interest (ROI). Phosphorylated α-synuclein (pS129-syn) spots were identified within the ROI mask from TUBB3-positive neurite. Total fluorescence intensity of pS129-syn spots per field was measured and normalized against total neurite length per field. The normalized total fluorescence intensity of pS129-syn spots was used as readout for α-synuclein aggregate formation.

### 2.7. Cell viability assay

Two-week differentiated DNs were treated with small molecules at required concentrations (Table S3) in Brainphys Neuronal medium supplemented with N2A and SM1 Supplements for two weeks. After treatment, cell viability tests were performed with CellTiter-Glo 2.0 Viability Assay kit (Promega, cat G9241) according to manufacturer’s instructions. The number of viable cells after treatment was measured by quantifying ATP according to luminescence readout from microplate reader (Molecular Devices, SpectraMax ID3).

## 3. RESULTS

### 3.1. Characterization of midbrain dopamine neurons from patient-derived iPSCs with an *SNCA* triplication mutation

Building on our earlier study, in which synuclein aggregation in midbrain organoids was highest when the *SNCA* triplication mutation was present, we chose this line and the respective matching isogenic controls for the work in this study [25]. These included an *SNCA* triplication line (*SNCA*-Tri), in addition to the isogenic control and *SNCA* KO lines. To assess the quality of the DA NPCs and DNs derived from each of the *SNCA* Tri, Isog Ctrl and the *SNCA* KO iPSC lines (see the schematic representation of the protocol (Fig. 1A), we performed IF staining on DA NPCs (Fig. 1C) and two-week DNs (Fig. 1B and 1D). Images of DA NPCs demonstrated that cells were positive for nestin, a type VI intermediate filament protein expressed on neural progenitor cells, SOX1, a transcription factor functioning primarily in neurogenesis, and LMX1A and FOXA2, markers that are expressed by floor plate progenitors (Fig. 1C). Staining of two-week differentiated DNs showed that >80% cells expressed neuron-specific class III-beta-tubulin (TUBB3) and microtubule-associated protein 2 (MAP2). Moreover, most of the two-week differentiated cells also stained positive for midbrain DN markers tyrosine hydroxylase (TH), FOXA2, Nurr1 and G protein-activated inward rectifier potassium channel 2 (GIRK2). Scoring the relative abundance of neurons with positive staining of different markers indicated that there were no significant differences between the three cell lines (Fig. 1C and 1D). To further characterize the progression of midbrain DN differentiation, gene expression profiles were evaluated at different stages (iPSC, DA NPC and two-week differentiated DN) using quantitative PCR (qPCR) (Fig. 1E). The expression of the key transcription factor for stem cell pluripotency, POU class 5 homeobox 1 (OCT3/4), decreased during the progression of differentiation.

**Figure 1.**
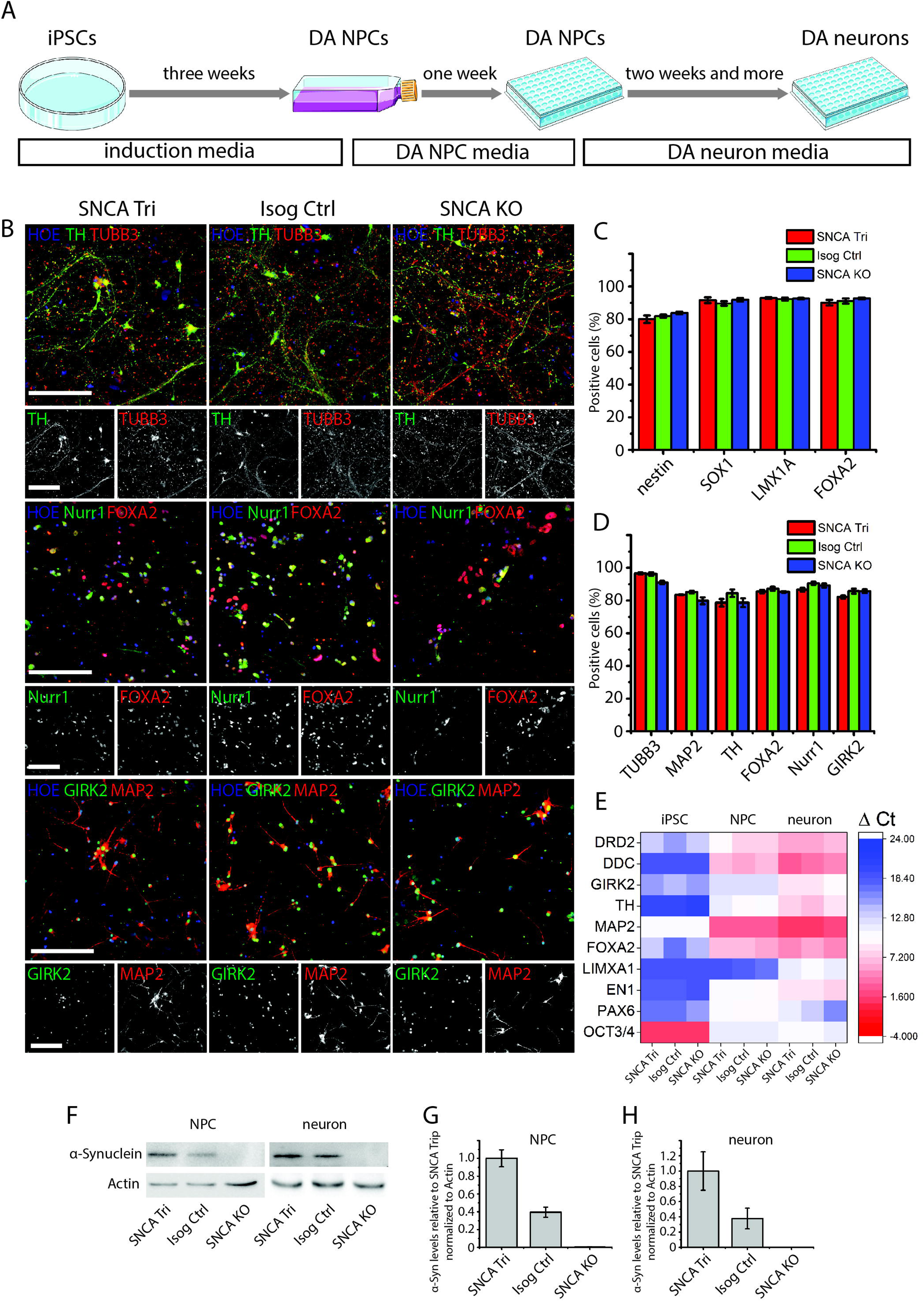
Generation and characterization of ventral midbrain neural progenitor cells and dopaminergic neurons from SNCA lines. (A) Schematic diagram of procedure to generate iPSC-derived midbrain neural progenitor cells and dopaminergic neurons. (B) Representative immunofluorescence images of staining for dopaminergic neuronal markers TH, TUBB3, Nurr1, FOXA2, GIRK2 and MAP2 in DA neurons after two-week differentiation from the SNCA-Tri line, its isogenic control line and SNCA KO line midbrain NPCs; bars = 150 μm. (C) Quantification of marker expression in DA NPC cultures labeled with antibodies for midbrain neural progenitor cell (NPC) markers Nestin, SOX1, FOXA2 and LMX1A and counter-stained with Hoechst 33342 (HOE); The data show the mean ± SEM; N = 12, from three independent experiments. (D) Quantification of marker expression in two-week old differentiated DA neuronal cultures; The data show the mean ± SEM; N = 8, from two independent experiments. (E) Heatmap of qPCR analysis showing gene expression level changes for markers of cell state during neuronal differentiation. (F-H) Western blot analysis of α-syn protein levels normalized to actin in midbrain NPCs (G) and two week differentiated DNs. (H) Quantification by densitometry (arbitrary units) (N = 3, mean ± SEM)

The expression of floor plate progenitor markers EN1, LMXA1 and FOXA2 increased during the midbrain differentiation, while the expression of PAX6, a transcription factor involved in neuroectoderm development, was increased in the DA NPC stage yet decreased during the process of neuronal differentiation. Neuronal markers MAP2 and midbrain dopaminergic markers TH, GIRK2, DDC and DRD2 were highly enriched in the neuronal stage, providing further evidence of the midbrain nature of the DNs generated. To confirm that α-syn levels differed across cell-lines due to differences in the *SNCA* copy number, α-syn its protein levels were assessed by immunoblotting lysates obtained from DA NPCs and two-week old DNs (Fig. 1F-H). The α-syn protein levels were significantly elevated in the *SNCA* Tri compared with Isog Ctrl, while α-syn proteins were undetectable in both *SNCA* KO DA NPC and DN samples. Taken together, our findings indicate that we successfully generated a set of DNs with different α-syn protein levels that also display cellular characteristics consistent with midbrain DN identity. Having established the expected characteristics of these cell lines, we employed them in the development of the seeding assay, as described in the following section.

### 3.2. α-synuclein PFFs elicit the formation of phosphorylated synuclein aggregates in iPSC-derived dopamine neurons

In a previous study [9], we showed that PFFs are rapidly taken up by DNs, leading us to test whether the addition of PFFs for a prolonged period would act as a seed for the formation of α-syn aggregates. Human α-syn PFFs were added to two-week old DNs derived from each of the three iPSCs lines (Fig. 2A). DNs were left for 2 weeks in the presence of PFFs, fixed and stained to assess if pS129-syn positive aggregates formed (Fig. 2B). Consistent with previous studies with primary cultured rat and mouse neurons, IF images confirmed that *SNCA* Tri and Isog Ctrl DNs contained aggregates that stained positive for pS129-syn 14 days after the addition of α-syn PFFs. The highest pS129-Syn signals were observed in the *SNCA* Tri DNs compared with Isog Ctrl DNs after 14-day PFF treatment (Fig. 2B). In contrast, the pS129-Syn signal was undetectable in *SNCA* KO neurons, in which endogenous synuclein expression is absent. PFFs were needed for the pS129-syn signal to be present, as in DNs derived from the *SNCA* Tri minimal signal was detected without PFF treatment (Fig. 2B).

**Figure 2.**
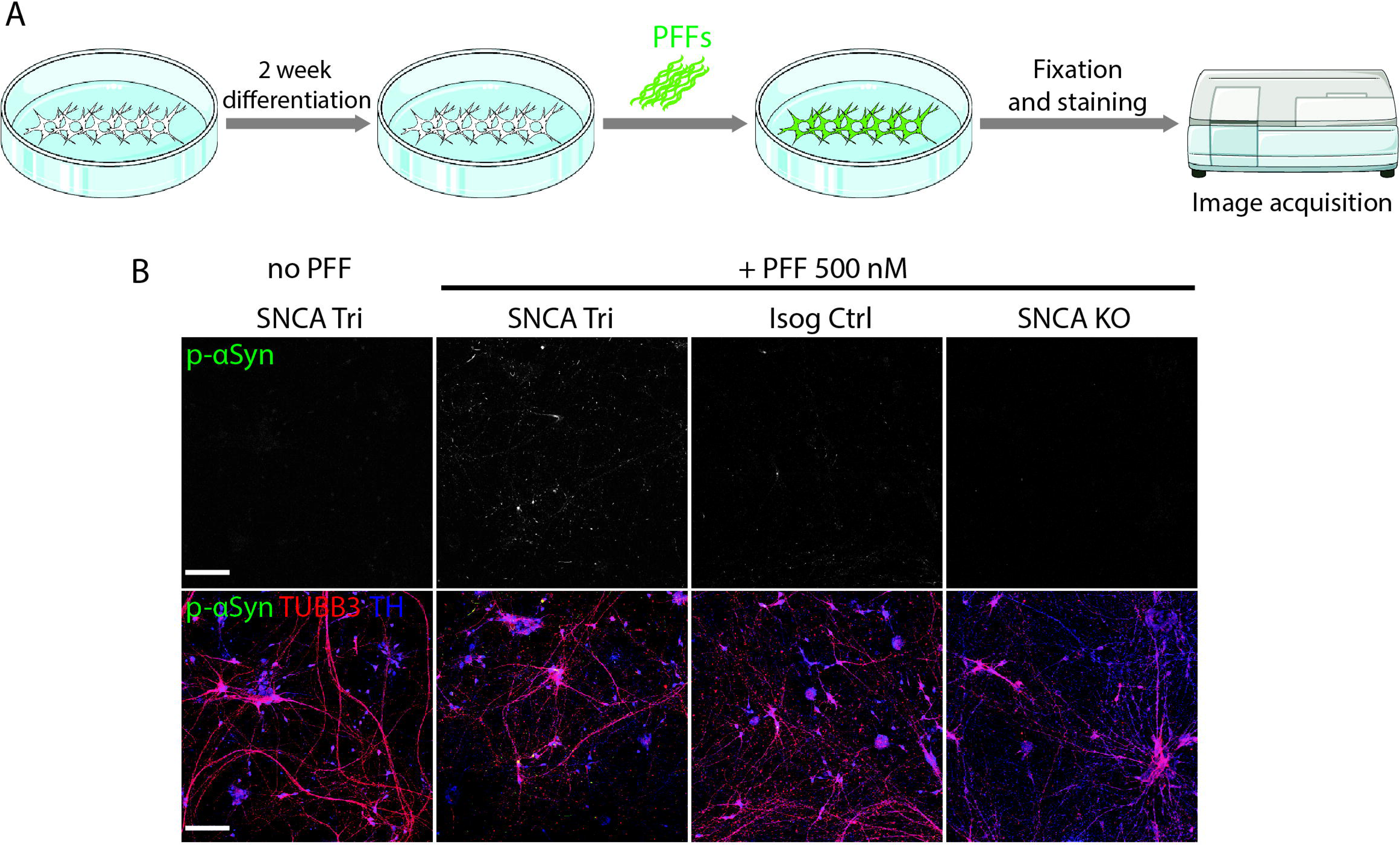
α-Syn PFFs trigger the formation of α-Syn aggregates in iPSC-derived dopaminergic neurons. (A) Overview of α-Syn aggregate formation experiments in which human DNs are differentiated from familial PD patient iPSCs with a triplication of the α-Syn gene (SNCA Tri) locus. After two-week differentiation, DNs are treated with 500 nM α-Syn PFFs. (B) Two weeks after PFF treatment, DNs are fixed and processed for immunocytochemistry staining with antibodies towards pS129-Syn and TUBB3; bars = 100 μm.

To characterize these aggregates in greater detail, higher magnification confocal images were acquired in the *SNCA* Tri DNs treated with PFFs for 14 days. Interestingly, filamentous-like pS129-Syn positive aggregates were observed in the neurites and dense inclusions were also observed in some neuronal cell bodies in the vicinity of the nucleus (Fig. 3A). Next, we sought to determine to what extent this model recapitulates the formation of the Lewy bodies in the brains of patients with synucleinopathy. Ubiquitin (Ub) [31] and SQSTM1/p62 [32] are two well-established LB markers. Our IF and confocal microscopy analysis showed that both filamentous-like neuritic inclusions and dense aggregates in the cell bodies were positive for both Ub (Fig. 3B) and p62 (Fig. 3C). Furthermore, we also observed markers for membranous organelles including lysosomal structures (Lamp1 positive staining, Fig. 3D), mitochondria (TOM20 positive staining, Fig. 3E) and Golgi structures (GM130 positive staining, Fig. 3F), predominantly in the dense aggregates in the vicinity of the nucleus. Our results demonstrated that the aggregates induced by PFFs in the hiPSC-derived human DNs shared certain morphological and biochemical properties similar to the LB observed in patient brains [33].

**Figure 3.**
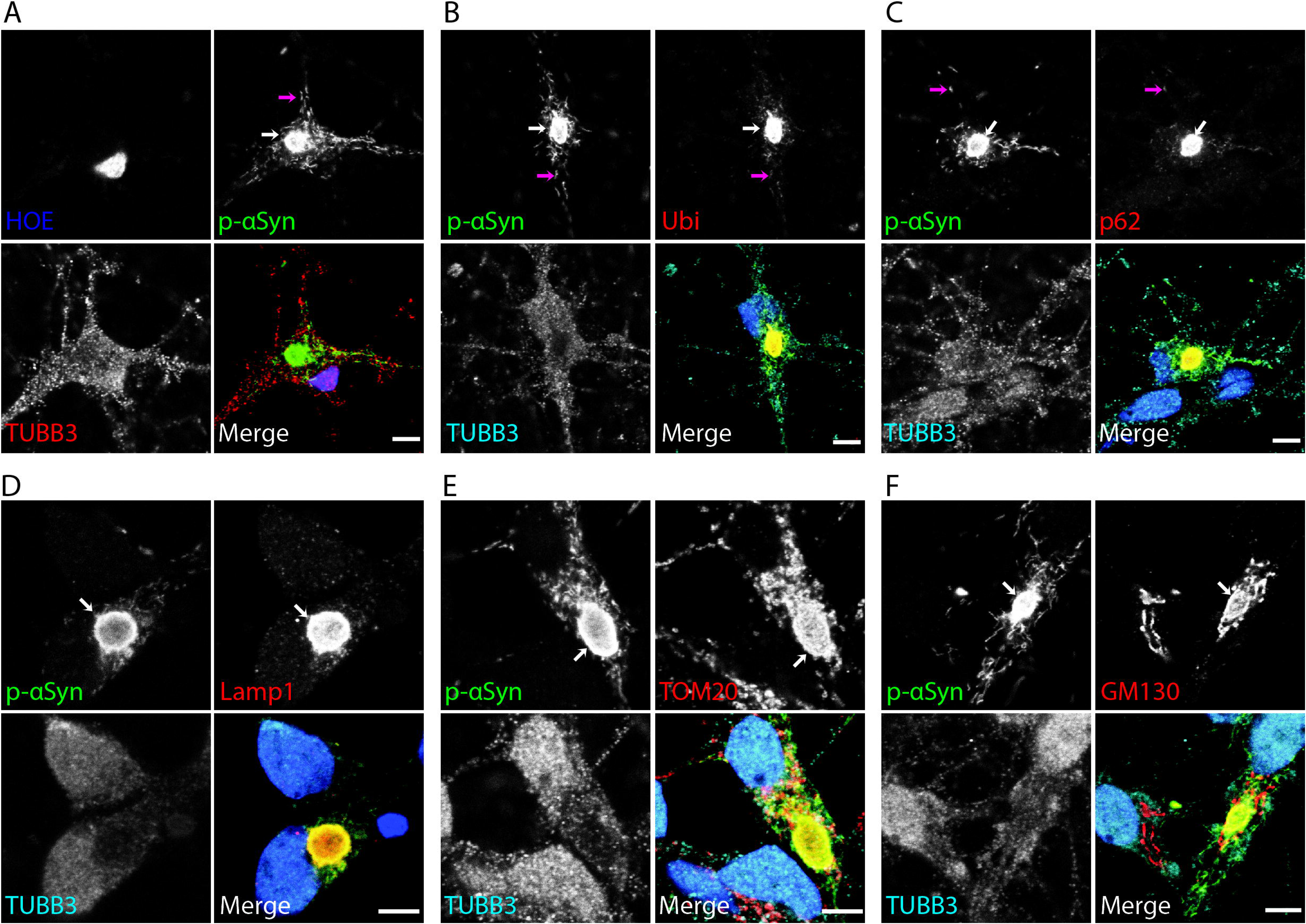
α-Syn PFFs trigger the formation of α-Syn aggregates in iPSC-derived dopaminergic neurons. α-Syn aggregates in PFF-treated DNs with the SNCA triplication mutation. Aggregates were detected by immunocytochemistry staining using either pS129-Syn in combination with TUBB3 (A),Ub (B) or with p62 (C). Bars = 5 μm. Aggregates were stained for pS129-Syn in combination with organelle-specific markers LAMP1 (late endosomes and lysosomes) (D), TOM20 (mitochondria) (E) or GM130 (Golgi) (F). DNs were counterstained with TUBB3 antibody and nucleus with HOE. Bars=5 μm

### 3.3. High-throughput imaging and analysis of the formation of α-synuclein aggregates

Next we implemented a scalable high content imaging assay workflow that could be used for testing small molecules and other agents. With this in mind, we plated *SNCA* Tri DA NPCs in 96-well microplates optimized for high-throughput imaging. After two-week differentiation, DNs were treated with PFFs (14 days for most conditions) before fixation and IF staining with Hoechst, TUBB3 and pS129-Syn. All the images were acquired and analyzed on a High-Content imaging Platform (Fig. 4A). With images acquired automatically from DNs treated with PFFs for 14 days, we observed a significant increase of pS129-Syn fluorescent signals compared with DNs treated with α-syn monomers (Fig. 4B). To quantify the level of PS129-Syn fluorescence, we set up an image analysis pipeline. In the analysis protocol, a mask was created to delineate the TUBB3-positive neurites (red channel) and a second mask was created around the identified spots of pS129-Syn (Green channel) (Fig. 4C). Next, total fluorescence intensity of pS129-Syn was quantified and normalized against total neurite length for each sample. Normalized total fluorescence intensity of pS129-Syn was used as readout for α-synuclein aggregate formation. To test the accuracy and efficiency of the imaging and data analysis pipeline, we treated *SNCA* Tri DNs with PFFs at different concentrations or with monomers. After fixation, some wells were incubated with Lambda protein phosphatase prior to the IF staining. We observed a significant increase in α-synuclein aggregation readouts (i.e., pS129-Syn intensity normalized with total neurite length) in the samples treated with PFFs in contrast with samples treated with monomers; and pS129-Syn intensity increased with the increasing concentration of PFFs added to the cultures. However, pS129-Syn signals were undetectable in the samples treated with Lambda protein phosphatase (Fig. 4D, 4E and 4F), showing that the antibody recognized specifically phosphorylated form α-syn. Thus, these findings demonstrated that the pipeline we developed to quantify the formation of α-syn aggregates in DNs is robust enough for high-throughput imaging and analysis.

**Figure 4.**
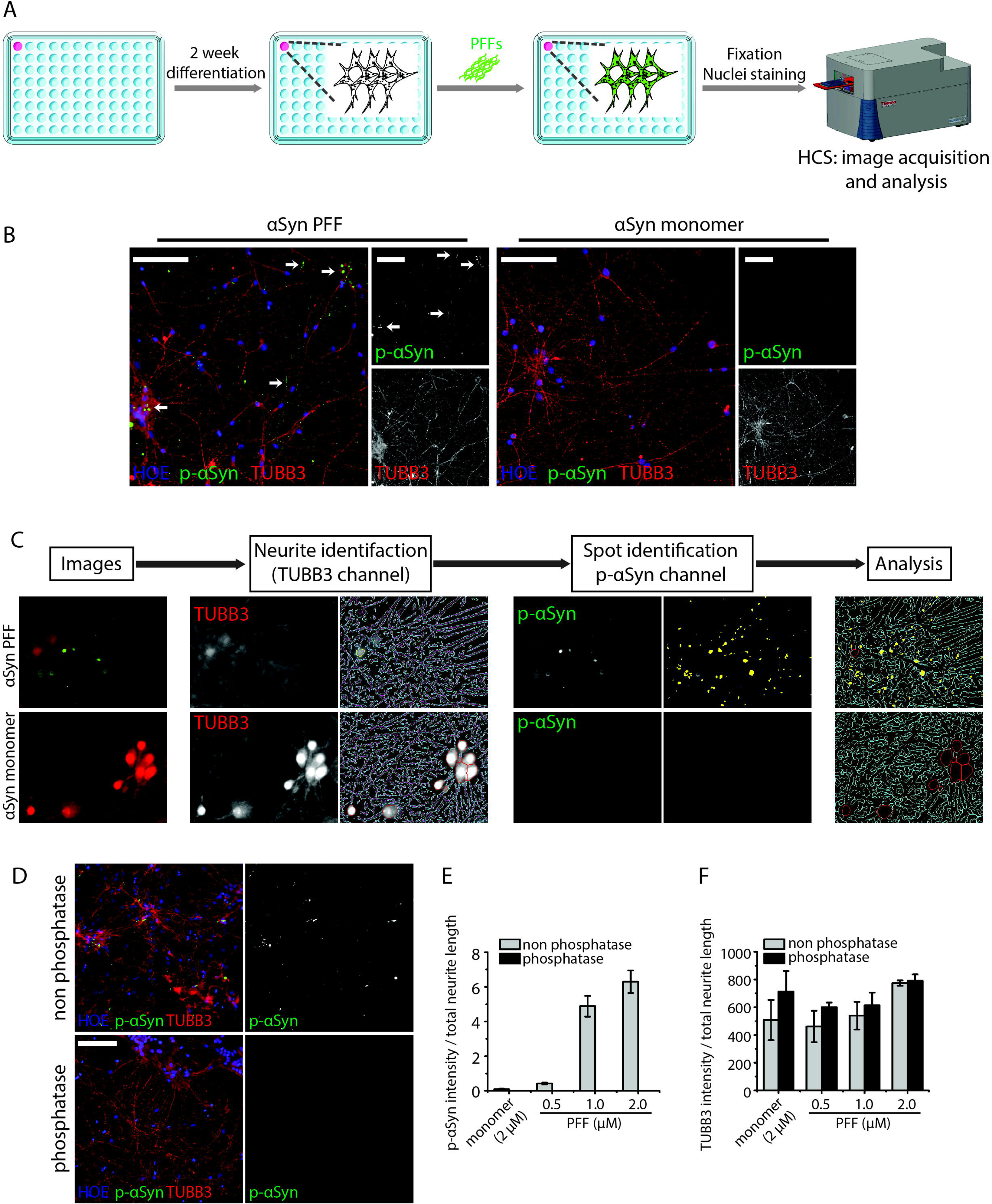
High-throughput imaging and analyzing the formation of α-Syn aggregates in iPSC-derived dopaminergic neurons. (A) Overview of pS129-syn aggregate formation assay set-up. (B) Images of phosphorylated synuclein aggregates in SNCA Tri DNs. Two week old DNs are treated with 500 nM α-Syn PFFs or 500 nM α-Syn monomers.Two weeks after the treatment, DNs are fixed and stained for pS129-Syn and TUBB3. Bars = 100 μm. (C) Quantification of pS129-Syn aggregate formation. Total neurite area is identified and a mask for the neurite area is created from the TUBB3 channel image. Spots of pS129-Syn aggregates are identified in the pS129-Syn channel and analyzed within the TUBB3 neurite mask. (D) Confirmation of phosphorylated signal in DNs. DNs from the SNCA Tri line are differentiated for two weeks and then treated with 1 μM PFF for two weeks. After fixation, DNs on coverslips are treated with or without Lambda Protein Phosphatase prior to staining for pS129-Syn and TUBB3. bar = 100 μm. (E and F), Quantification of total pS129-Syn signal (E) and TUBB3 fluorescent signal (F) in SNCA_tri DNs; the data represents the mean±s.e.m. N=3 replicates.

### 3.4. α-synuclein aggregate formation and compound testing in PD patient and control dopamine neurons

In order to validate our α-syn aggregate formation assay and to confirm that pS129-Syn is a valid measurement to quantify α-syn aggregates in DNs from PD and control cell lines, we treated *SNCA* Tri and Isog Ctrl DNs with PFFs at different concentrations for 14 days. As shown in Fig. 5A, PFFs induced α-syn aggregate formation in a concentration-dependent manner in both cell lines; however, the signal for α-syn aggregates was significantly higher in the *SNCA* Tri DNs compared with Isog Ctrl DNs treated with PFFs at same concentration. In *SNCA* Tri DNs, treatment with 250 nM PFFs triggered a significant increase in the α-syn aggregate signal relative to untreated *SNCA* Tri DNs or *SNCA* Tri DNs treated with α-syn monomers. For the Isog Ctrl DA, a PFF concentration of 500 nM was necessary to trigger a significant increase in α-syn aggregation. For the robustness of the assay, we chose 500 nM PFFs as our treatment condition and performed a temporal analysis of α-syn aggregate formation at Day 4 (Fig. 5B), Day 7 (Fig. 5C), Day 14 (Fig. 5D) and Day 28 (Fig. 5E). We observed that the PFFs increased pS129-Syn signal in a time-dependent manner. We started to obtain robust α-syn aggregation readouts for *SNCA* Tri DNs 7 days after PFF treatment; whereas for the Isog Ctrl DNs, a significant increase in α-syn aggregation signals was observed 14 days after PFF treatment (Fig. 5F); with no detectable increase in α-syn aggregation with *SNCA* KO DNs. Such findings in the KO lines support the idea that endogenous synuclein is needed for aggregates to form. Together, these findings confirm that pS129-Syn signal is dependent on endogenous α-syn and that our α-syn aggregate formation assay with the high-throughput imaging and analyzing method is sensitive enough to indirectly assess copy number variants of the SNCA gene.

**Figure 5.**
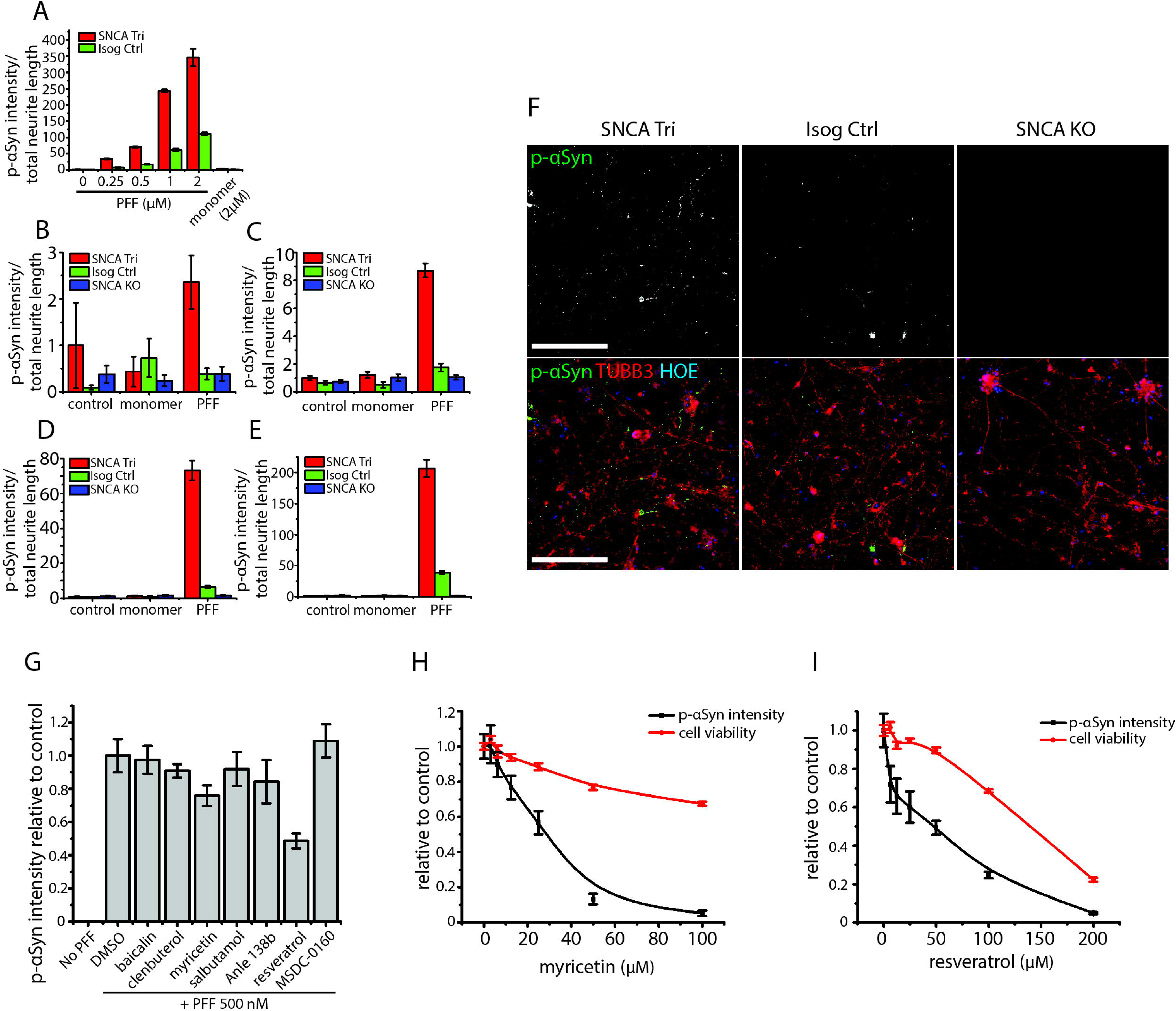
αSyn aggregate formation assay optimization and compound tests in iPSC-derived dopaminergic neurons. (A) Dose-dependent increased of pS129-Syn aggregates in α-Syn PFF treated DNs from SNCA Tri and Isogenic control lines; the data shows the mean ± SEM, N = 5. (B, C, D and E) Temporal analysis of phosphorylated α-Syn aggregates formed at Day 4 (B), at D7 (C), at D14 (D) and at D28 (E) post-α-Syn PFF treatment; the data shows the mean ± SEM; N = 8, from two independent experiments. (F) Representative images of four weeks DNs following a α-Syn PFF treatment of 14 days. The IF shows that DNs from SNCA Tri line present higher number of puncta of pS129-Syn aggregates than its isogenic control line. No pS129-Syn aggregate where detectable in the SNCA KO line; bars = 100 μm. (G) Phosphorylated α-Syn signals in SNCA Tri DNs treated with different compounds compared to DMSO-alone treatment.; the data shows the mean ± SEM N = 6, from two independent experiments. (H and I) Dose-response curves of myricetin (H) and resveratrol (I) in cell viability assays (red line) and αSyn aggregate formation assays (black line) on DNs derived from the SNCA Tri iPSC line; the data shows the mean ± SEM; N = 8, from two independent experiments.

As proof of principle, we performed this assay with a small set of compounds which were previously reported to affect α-syn aggregate formation in other experimental models [34–39]. First, we treated two-week differentiated SNCA Tri DNs with 500 nM PFFs along with DMSO as control or with one concentration of baicalin, clenbuterol, myricetin, salbutamol, Anle 138b, resveratrol or MSDC-0160 (see Table S3 for the concentrations). After 14 days, DNs were subjected to fixation, IF, high-throughput imaging and analysis. We observed a robust increase in the pS129-Syn signals in PFF-treated DNs in contrast to non-treated controls. Interestingly, two of the compounds tested, myricetin and resveratrol elicited a significant decrease in the level of pS129-Syn signal compared to DMSO controls: (Fig. 5G). Next, we tested dosage responses (Table S3) of myricetin and resveratrol in the α-Syn aggregate formation assay. As shown in Fig. 5H and 5I, both compounds decreased the α-syn aggregation readout in a dose-dependent manner. To determine if the compounds were eliciting toxicity, we assessed the viability of the DNs following the addition of increasing concentration of the myricetin and resveratrol. As shown in Fig. 5H and 5I, both compounds decreased viability, in a dose-dependent manner. However, low doses of myricetin (12.5 and 25 μM) and resveratrol (6.25 to 50 µM) could decrease pS129-Syn aggregation while maintaining a viability rate above 85% relative to untreated conditions. Taken together, we demonstrated that the application of synuclein fibrils to elicit α-syn aggregate formation in iPSC-derived DNs can be used to test for small molecules, leading to the validation of two compounds previously shown to reduce synuclein aggregate formation.

## 4. DISCUSSION

In this paper, we report for the first time a scalable screening assay monitoring the formation of α-syn aggregates in human DNs with the potential for use in assessing novel therapeutics developed for synucleinopathies. Following the protocol developed in our laboratory, we generated ventral midbrain DNs from an iPSC-line derived from a patient with an *SNCA* gene triplication (*SNCA* Tri) along with an isogenic control iPSC line (Isog Ctrl) and a *SNCA* KO iPSC line. We did not observe any significant difference among these DNs during the process of induction and differentiation other than the endogenous α-syn protein levels, which implies that α-syn might not be functionally involved in the DN development. However, with the addition of α-syn PFFs to the cultures, α-syn aggregation differed among the DNs from the three iPSC lines. While both *SNCA* Tri and Isog Ctrl DNs formed α-syn aggregates following exogenous PFFs, in a dose and time dependent manner, DNs with *SNCA* triplication presented with levels of detectable α-syn aggregates following a shorter PFF treatment time compared with Isog Ctrl DNs. Moreover, after the same PFF treatment period, *SNCA* Tri DNs also presented with higher levels of α-syn aggregates relative to the Isog Ctrl. In the *SNCA* KO, we could not detect a pS129-Syn signal at any of the doses or time-points following PFF treatment, in support of the idea that endogenous synuclein is required for aggregates to form. Altogether, our results demonstrate that exogenous PFFs recruit endogenous α-syn to form aggregates, and that increased level of endogenous cellular α-syn accelerates the rate and extent of aggregate formation.

In our study, we also assessed the properties of the α-syn aggregates in the human iPSC-derived DNs. Posttranslational modifications of α-Syn, particularly phosphorylation at serine 129 (pS129) are common in synucleinopathies especially in PD. More than 90% of α-syn in LBs from patients with sporadic and genetic forms of PD is phosphorylated at S129 [40]. Extensive in vitro studies demonstrated that pS129, along with other PTMs and protein misfolding, can alter the characteristics of α-syn such as propensity of aggregation, neurotoxicity, inclusion formation and cell-to-cell transmission [41–44]. Phosphorylated α-syn is widely used as a biomarker clinically as well as in modeling synucleinopathies experimentally [45]. In our study, we chose phosphorylated α-syn at serine 129 (pS129-Syn) as our major readout to develop the aggregation formation assay. We observed filamentous-like pS129-Syn positive aggregates in the neurites and dense inclusions in some neuronal cell bodies in the vicinity of the nucleus in *SNCA* Tri DNs treated with PFFs for 14 days, which is consistent with previous observations in primary cultured rat and mouse cortical neurons [4]. As Ub and p62/sequestosome 1 have also been identified as common components of LBs in PD patient brains [32], we have also looked at these two markers. We demonstrated that Ub and p62 were highly enriched in the pS129-Syn positive dense inclusions in the neuronal cell bodies. In contrast, we only detected a very low signal for Ub and p62 in the filamentous-like pS129-Syn positive aggregates in the neurites. Thus, similar to observations with LBs from PD patient midbrain sections, our results indicate the possibility of dissimilarities in the formation of aggregates in the cell bodies versus the neurites [4, 32]. The differential localization of Ub and p62 suggests that they might play a role in the early neurofibrillary pathogenesis. As such, differential staining patterns have also been observed in the Tau aggregation process [46]. From ultrastructural analysis of LBs on postmortem human brain tissue from PD and other synucleinopathies, a crowded environment of membranes and organelles are identified in the LBs [33]. We observed the recruitment of membranous organelles markers (lysosomes, mitochondria and Golgi) in pS129-Syn positive aggregates, especially in the dense inclusions in the vicinity of the nucleus. Our observations are supported by Bayati et al (2024) ^[10]^ who showed by electron microscopy ultrastructural analysis that the aggregates induced by PFFs in the hiPSC-derived human DNs shared certain morphological and biochemical properties of the *bona fide* LB observed in patient brains.

To develop the model of α-syn aggregation in DNs into a scalable screening assay, we built a high-throughput imaging and analysis pipeline. In our current study, images were automatically acquired on a High-Content Screening Platform and analyzed with a pipeline based on neuronal profiling. In our experiments, we observed a significant increase in α-syn aggregation in the samples treated with PFFs compared to those treated with monomers. Moreover, we also demonstrated that the PFFs induced endogenous α-syn to form phosphorylated aggregates in a time-dependent and concentration-dependent manner. In addition, we performed this assay with a set of compounds previously reported to affect α-syn aggregate formation in other experimental models as a proof of principle. In our screening, two compounds, myricetin and resveratrol, showed significant inhibition of PFF-induced α-syn aggregate formation. Myricetin is a flavonoid of subclass flavonols. It has been reported that myricetin can interact with α-syn monomers and prevent the oligomerization and reduce synaptic toxicity in mouse models [47, 48]. Resveratrol is a polyphenolic compound derived from grapes and red wine, and has been shown to inhibit α-syn aggregation and promote α-syn autophagic degradation in MPTP-treated mice [49, 50]. Our subsequent dose response tests with these two compounds suggested that myricetin and resveratrol might represent potential therapeutic agents for synucleinopathies and highlight how the assay can be applied towards assessing candidate therapeutics.

For this study, we focused on α-syn aggregate formation in hiPSC-derived DNs. However, with some minor adaptations, this assay and its imaging and analyzing pipeline could be used in other High-Content Screening Platforms and analysis software. Moreover, our automated α-syn aggregation assay can be used to study other α-syn PTMs that have been identified in postmortem brains of PD patients, such as acetylation at the N-terminus and lysine residues, nitrations and truncations as well as phosphorylation at other amino acids [51–53], dependent on the availability of high quality, validated antibodies for immunofluorescence readouts.

We acknowledge that there are several limitations in our assay. Here, we only focus on aggregation in a single cell type. Even though degeneration of DNs in the substantia nigra par compacta is the classical hallmark associated with the symptoms of motor impairment in PD, evidence is increasing that non-neuronal cells, such as glial cells, also play a crucial role in PD and other synucleinopathies. Under physiological conditions, neurons in the brain communicate with microglia and astrocytes to maintain parenchymal structure and function; but in α-syn related pathological states, astrocytes and microglia play a major role in the degradation of α-syn aggregates and α-syn in the pathogenic form (oligomer, aggregates) can trigger the gliosis of these cells, leading to the release of neurotoxic chemokines and cytokines [10, 54–58]. Hence, we see benefit in adapting this assay further towards monitoring α-syn aggregate formation in co-culture systems with different glial cell types. In addition, for our current assay, we focused on aggregate formation in which we chose phosphorylated α-syn as our major readout. However, we didn’t observe DN degeneration related phenotypes (such as neurite fragmentation, cell death) in our experimental condition. Possible reasons could be insufficient PFF exposure time, inadequate DN maturity or lack of glial cells. Hence, further study is needed to examine whether more toxic PFF species can be used.

Taken together, we have developed an automated α-syn seeding assay to quantify the extent of α-syn aggregation in hiPSC-derived DNs from patients with PD. Our α-syn seeding assay can be used to test the capacity of compounds to impede or reduce α-syn aggregation and the workflow can be easily adapted to study other PTMs or other synucleinopathy-relevant cell types in further studies.

## Author Contributions

Conceptualization, C.H.; Methodology, C.H., E.N.-R.,F.B., W.L., I.S. and E.D.C.P,; Software, A.K.F.-F. and W.E.R.; Validation, C.H., F.B. and T.M.D.; Formal analysis, C.H. and F.B.; Investigation, C.H., E.N.-R., F.B., C.X.-Q.C., A.A., L.V.; Data curation, C.H. and E. N.-R.; Writing—first draft preparation, C.H.; Writing—review and editing, C.H., F.B. and T.M.D.; Visualization, C.H.; Supervision, T.M.D.; Project administration, C.H. and T.M.D.; Funding acquisition, L.K.B., E.A.F and T.M.D. All authors have read and agreed to the published version of the manuscript.

## Acknowledgements

We thank Genevieve Dorval for administrative assistance, the Neuro’s Early Drug Discovery Unit’s Microscopy Core Facility and the Neuro’s microscopy Core Facility.

