## Supplemental Information for "An automated workflow for quantifying the formation of synuclein aggregates in human dopaminergic neurons"

**Supplement Information**

**Figure S1**

**Production of recombinant α-syn monomers and preformed fibrils (PFFs).**

(**A**) Overview of PFFs production. (**B**) polyacrylamide SDS-PAGE gel of BL-21 cell lysate expression recombinant GST- α-syn (cell lysate), GST-tagged α-syn (purified GST-Syn), protein sample post 3C protease cleavage of the GST tag (protease digestion) and final purified untagged α-syn. (**C**) Representative EM microphotographs of non-sonicated α-syn PFFs and α-syn PFFs sonicated for 30 seconds. Bars = 200 nm. (**D**) Representative size distribution of fibrils within sonicated α-syn PFFs assessed through quantification of particle size by dynamic light scattering.

**Table S1. Antibodies used for immunofluorescence microscopy.**

| Antibodies | Source | Cat# | RRID | Dilution |
|---|---|---|---|---|
| Nestin | Abcam | AB92391 | AB_10561437 | 1/150 |
| SOX1 | R&D system | AF3369 | AB_2239879 | 1/100 |
| LMX1A | Millipore | AB10533 | AB_10805970 | 1/500 |
| FOXA2 | R&D system | AF2400 | AB_2294104 | 1/200 |
| TUBB3 | Millipore | AB9354 | AB_570918 | 1/3000 |
| MAP2 | Encor Biotech | CPCA-MAP2 | AB_2138173 | 1/500 |
| TH | Peel -Freez | P40101-150 | AB_2617184 | 1/400 |
| Nurr1 | Santa Cruz | SC-990 | AB_2630633 | 1/100 |
| GIRK2 | Millipore | AB5200 | AB_91747 | 1/200 |
| P-α-synuclein | Abcam | AB51253 | AB_869973 | 1/2000 |
| Ubiquitin | Avacta | AVA00100 | N/A | 1/200 |
| p62 | Cell Signaling | Q13501 | AB_2800125 | 1/400 |
| TOM20 | Santa Cruz | SC-17764 | AB_628381 | 1/400 |
| GM130 | BD | 610823 | AB_398142 | 1/400 |
| Dylight 488 rabbit | Abcam | ab96891 | AB_10679664 | 1/500 |
| AlexaFluor647 chicken | Jackson Immuno Research | 703-605-155 | AB_2340379 | 1/500 |
| AlexaFluor647 goat | Invitrogen | A21447 | AB_141844 | 1/500 |
| Dylight 550 mouse | Abcam | ab96876 | AB_10679663 | 1/500 |

**Table S2. List of TaqMan probes and primer sets (Applied Biosystems).**

| Gene ID | Gene Name | Accession Number | Assay ID |
|---|---|---|---|
| OCT3/4 | POU class 5 homeobox 1 | NM_001173531.2 | Hs04260367_gH |
| PAX6 | paired box 6 | NM_001127612.1 | Hs01088114_m1 |
| EN1 | Engrailed Homeobox 1 | NM_001426.3 | Hs00154977_m1 |
| LMXA1 | LIM homeobox transcription factor 1 alpha | NM_001174069.1 | Hs00898455_m1 |
| FOXA2 | Forkhead Box A2 | NM_021784.4 | Hs00232764_m1 |
| MAP2 | Microtubule associated protein 2 | NM_001039538.1 | Hs00258900_m1 |
| TH | Tyrosine Hydroxylase | NM_000360.3 | Hs00165941_m1 |
| GIRK2 | potassium voltage-gated channel subfamily J member 6 | NM_002240.4 | Hs01040524_m1 |
| DDC | Dopa Decarboxylase | NM_001082971.2 | Hs001105048_m1 |
| DRD2 | dopamine receptor D2 | NM_000795.3 | Hs00241436_m1 |

**Table S3. Small-molecules tested**

| Compounds | Supplier | Catalog number | Seeding assay Doses (µM) | Viability assay doses (µM) | References |
|---|---|---|---|---|---|
| Baicalin | Cayman | 19843 | 2.5 | - | [1] |
| Clenbuterol | Cayman | 14985 | 5 | - | [2] |
| Myricetin | Cayman | 10012600 | 5 | 0, 3.125, 6.25, 12.5, 25, 50, 100 | [3] |
| Salbutamol | Cayman | 21003 | 5 | 0, 25, 50, 100, 200, 400, 800 | [2] |
| Anle138b | Cayman | 34259 | 5 | - | [4] |
| Resveratrol | Cayman | 70675 | 5 | 0, 6.25, 12.5, 25, 50, 100, 200 | [5] |
| MSDC-0160 | Cayman | 71748 | 2.5 | - | [6] |
