## Supplementary figures and images for "An automated workflow for quantifying the formation of synuclein aggregates in human dopaminergic neurons"

### Supplemental Figure S1

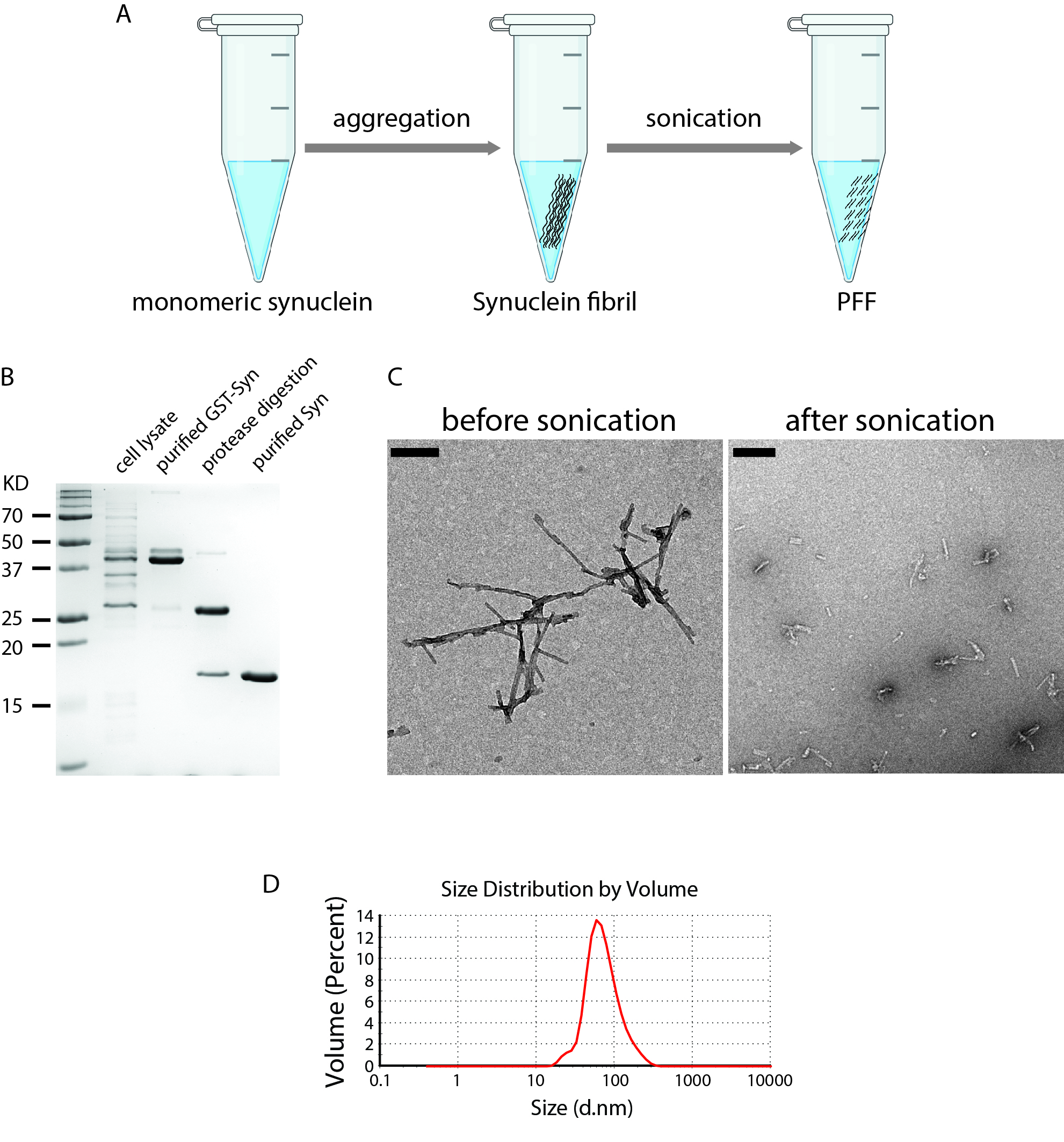
